# mzQuality: A tool for quality monitoring and reporting of targeted mass spectrometry measurements

**DOI:** 10.1101/2025.01.22.633547

**Authors:** Marielle van der Peet, Pascal Maas, Agnieszka Wegrzyn, Lieke Lamont, Ronan Fleming, Amy Harms, Thomas Hankemeier, Alida Kindt

## Abstract

Analyzing metabolites using mass spectrometry can offer valuable insight into an individual’s health or disease status. However, various sources of experimental variation can affect the data, making robust quality control essential. In this context, we introduce mzQuality, a user-friendly software tool designed to evaluate and correct technical variations in mass spectrometry-based metabolomics data. MzQuality offers key quality control features, such as batch correction, outlier identification, and analysis of signal-to-noise ratios. It supports any peak-integrated processed data independent of vendor software and does not require the user to have any programming skills. We demonstrate the functionality of mzQuality with a data set of 419 samples measured across six batches, in which mzQuality effectively minimized experimental variation, ensuring the data’s readiness for statistical analysis and biological interpretation. With customizable settings, mzQuality can be seamlessly integrated into research workflows to produce more accurate and reproducible metabolomics data.

## Introduction

Metabolites are influenced by both endogenous factors and an individual’s immediate environment, such as diet, living arrangements, or other, more general lifestyle factors. All of these metabolites are reflected in the metabolome^1^. Analyzing the metabolome, also known as metabolomics, can provide valuable information on an individual’s health status and help elucidate disease-related metabolic phenotypes. This can be of clinical interest when investigating disease progression, possible treatment outcomes, or even biomarkers for early detection and early intervention^2^.

Targeted Mass Spectrometry (MS)-based analytical techniques are often used to detect and quantify hundreds of metabolites simultaneously. Mass spectrometers are generally combined with separation techniques, such as liquid chromatography (LC), due to their high sensitivity and selectivity^3^. However, in practice, it can be challenging to acquire high-quality MS-based metabolomics data due to various sources of experimental variation. Sources of variation can be introduced during sample handling, sample preparation, and/or varying experimental conditions, such as differences in chromatographic mobile phases, column temperature, column pressure variation, instrumentation, instrumental drift, and even regular maintenance^4,5^. It can be challenging to keep these conditions stable in larger metabolomics studies where samples are divided, prepared, and measured over multiple subsets, termed batches, that are analyzed individually^6,7^.

All these experimental variations can collectively contribute to less reliable and reproducible results, which makes it harder to interpret the data properly and draw meaningful conclusions from the findings. To minimize variation as much as possible, it is important to have quality assurance (QA) guidelines and corresponding quality control (QC) practices to evaluate and correct data variability^8,9^. These guidelines should include protocols for the study design, sample collection and storage, sample preparation, and sample measurements^9^. An important part of QC practices is the use of QC samples, which can consist of long-term reference material (LQC) or short-term reference material (SQC). LQCs are used over extended periods and function as a reference over time to monitor the data quality throughout different studies SQCs, also named intra-study QC samples, often consist of pooled study samples that contain an average amount of metabolites representative of the study. Additional QC practices should include a systems suitability test (SST) to assess analytical stability, measurements of blanks to evaluate background signal, and the use of internal standards (IS) for normalization strategies^10^.

Even when variation is minimized as much as possible, it is still important to assess data thoroughly. Several open-source data processing tools, such as OpenMS^11^ and MZmine^12^, offer features like baseline noise filtering, retention time alignment, and a few quality metrics. However, these tools do not address the variations in sample measurements over extended periods. On the other hand, msQuality package by Naake et al, focuses more on quality metrics but does not automatically correct the identified issues^13^.

As the field of metabolomics continues to evolve, the integration of innovative software solutions becomes increasingly important to unlock the full potential of the field and contribute to more reliable findings. In this paper, we introduce mzQuality, a new metabolomics data quality assessment tool. Unlike other tools that process peaks, mzQuality distinguishes itself by providing a user-friendly interface for evaluating and correcting technical variations in the data without requiring extensive programming skills. Recognizing the importance of experimental design and procedural quality in metabolomic studies, we included guidelines as to how to implement mzQuality into a metabolomics workflow and optimize the software’s application^9^. MzQuality performs several important quality control steps, such as identifying sample outliers, calculating background signal, performing batch correction, detecting trends in total signal intensity for either metabolite or IS, and calculating metrics that highlight variation in metabolite/IS ratio are readily performed. In addition, compounds with a low signal-to-noise ratio are flagged for removal by the user. MzQuality uses repeat measurements of quality control samples to flag compounds that do not reach predefined thresholds as low quality. We defined the threshold at a relative standard deviation QC>30%, which follows the guidelines of the metabolomics quality assurance and quality control consortium^14^, and the background signal cannot be higher than 40%^15^. Furthermore, MzQuality allows users to adjust settings according to their preferences and analytical methods. How these assessment tasks are performed is showcased here using a large example data set included in GitHub: https://github.com/hankemeierlab/mzQuality, consisting of 419 study samples analyzed over six batches. Through this paper, we showcase how mzQuality identifies and corrects variations within the data, illustrating its functionality and effectiveness.

## Materials and methods

### Software information

MzQuality is distributed as an R package, for which version 4.0 or later is required. It is available through GitHub: https://github.com/hankemeierlab/mzQuality. For additional information on installation and usage, see the provided manual on GitHub.

### Example dataset

An example dataset obtained in our laboratory showcases the utility and effects of mzQuality. This data set comprises 6 random yet consecutively measured batches containing 419 EDTA plasma study samples originally part of the CoSTREAM consortium (https://costream.eu/index.html). Also included in the batches were different types of QC samples, blank samples, and calibration curves. The samples were prepared and measured according to the method described by Yang. et al (2024)^16^. Briefly, a targeted LC-MS method is operated in polarity switching mode where all metabolites and corresponding stable isotope labeled IS are analyzed in dynamic Multiple Reaction Monitoring (dMRM) mode. The data has been integrated using SCIEX OS (version 2.1.6.59781) and exported using the recommended data format.

### Results and discussion

Here, we discuss several steps needed for or performed by mzQuality: 1) the recommended batch design including the different sample types, 2) the data input format, 3) the calculations performed, 4) the generated output plots used for assessment, and 5) the report format. The example data set was used to showcase these steps.

### Batch design

Different sample groups or phenotypes should be randomly distributed across batches ensuring a balanced representation of different phenotypes within the study. If phenotypes are not balanced well, batch-related differences might introduce errors in downstream data analysis^17^.

The recommended order of sample injections begins with a system suitability test (SST), highlighted in orange, followed by one or more blank injections, as shown in Figure 1. SST samples are used to ensure the analytical system functions within predefined criteria to detect any potential contamination issues^8^.

**Figure 1:**
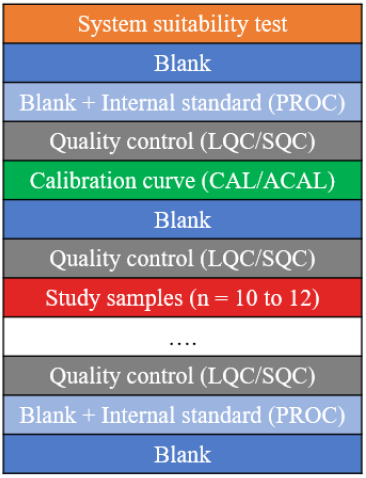
Batch design. The figure shows a summarized batch design, starting with a system suitability test. Quality control samples must be dispersed throughout the whole batch and at least every twelve injections. The quality control and study sample block can be repeated.

Blank samples are water samples prepared without the addition of IS. Following this, blank samples with added IS are injected (Procedure blank; PROC), these are highlighted in light blue. It is important to inject blank samples both at the beginning and the end of each batch, with a minimum of two injections. These samples can be used to assess carry-over and background signals in the measured data.

Although a calibration curve is not mandatory for the tool’s functionality, it can be incorporated into the batch design. Calibration curves can be made by adding an increasing concentration of analytical standard mix to water (ACAL) or biological matrix (CAL). While mzQuality does not use the calibration lines for corrections, they can calculate absolute concentrations and inform about the dynamic range in which samples are measured.

It is important to consider whether to use LQC, SQC, or a combination of both in the batch design, as these samples are used for batch correction and assessment of technical variation as assessed by the RSDqc per compound. If SQCs were generated from pooling study samples, then they provide an average signal over the study samples therefore allowing a direct comparison with the samples investigated with the biological samples. For example, if any technical drift were to occur, the impact on SQC samples would be most representative of the actual response in the study samples. This makes batch corrections based on SQC samples the most reliable^8^.

LQCs provide information on stability over time thus allowing us to monitor instrument stability and sensitivity beyond the duration of one study and correct for batch effects when a project was measured over a long period. When a compound is not detected in LQC or SQC samples, the compound will be assumed as not present and will not be reported by mzQuality.

It is recommended to have a minimum of four QC samples per batch and, depending on batch size, evenly spaced throughout the batch. We recommend one injection for every 15 study samples (highlighted in red). The injections of QC samples must be evenly distributed throughout the whole batch to check for instrument and extraction variance.

Replicates of study samples are treated independently, meaning no average of these samples is calculated or utilized for any analysis or correction. However, Rosner’s test is used on the IS of study samples to identify and flag potential misinjections and thus outliers. The complete batch design of the example set can be found in Supplementary Table 1.

### Data input and data validation

Chromatographic peak areas can be integrated using open-source or vendor software. The peak areas of compounds and IS, if applicable, need to be exported for use in mzQuality. We strongly recommend using a tab-separated format while exporting; an example can be found in GitHub: https://github.com/hankemeierlab/mzQuality.

Additionally, it is important that numbers and symbols are compatible with R. The input file should include the following columns with the specified headers (Table 1): aliquot, type, batch, compound, area, and datetime of the sample measurement. For quantitative or semi-quantitative analysis, the columns compound_is and area_is must also be included. Additional columns can be added for visualization, such as replicate number or retention time, however, this information is not used for corrections.

**Table 1:** Input file.

| Column name | Includes which info: |
| --- | --- |
| aliquot | Unique sample names |
| type | Sample type; SQC, LQC, BLANK, PROC, SAMPLE, CAL..., ACAL..., SST |
| batch | Batch number |
| compound | Compound name |
| area | The integrated peak area of the compound. These cells must have a numerical value or NA. |
| compound_is | Internal standard name |
| area_is | The integrated peak area of the internal standard. These cells must have a numerical value or NA. |
| datetime | The time the samples have been measured. The format of the date/time column should be year-month-day and hours:minutes:seconds, separated by a space. |

MzQuality performs a validation on the input file to ensure all necessary information is included and to detect any formatting errors. It checks for the presence of the required columns and verifies if a batch is assigned to all aliquots. If no batches are defined, all samples are assigned to batch 1.

During the validation process, mzQuality checks if the column “type” includes the sample type QC. Additionally, it verifies if there is an observation for every aliquot and every compound and inserts “NA” for any missing data. Any NA value is treated as an absent data point and will not be used for calculations. Finally, the injection order is checked based on the datetime column. If any of the necessary columns are absent mzQuality will not be able to run.

### Data processing

Once the data is imported and validated, mzQuality initiates the first steps of data evaluation to ensure data integrity.

#### IS normalization

The first step involves IS normalization, where for all samples ratios are calculated by dividing the compound area (Peak area_compound) by the corresponding IS area (Peak area_IS) as shown in equation 1.

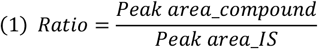

ISs can be predefined in the input file and should be selected based on class behavior and retention times. MzQuality also provides suggestions for alternative ISs, but the user must check the compatibility of the IS with the compound. These suggestions are based on calculations that evaluate all compound and IS combinations to determine which combination gives the best RSDqc after batch correction.

If no IS is available, mzQuality assumes an IS area of 1, meaning the area ratio, on which most of the calculations are based, is the same as the area.

#### Sample outlier determination

The next steps involve the identification of QC and study sample outliers using Rosner’s test. For QC samples, outliers are identified based on the median area/area_IS ratios, as these samples are assumed to be identical. While for study samples only the IS areas are used since they may vary biologically. Potential outliers are flagged by mzQuality and are automatically excluded from further calculations^18^. For study samples, the user needs to determine whether an outlier is biological or technical and whether it should be included in further analysis.

#### Batch correction

To correct for technical variation between individual batches by, e.g., sample preparation, mobile phases, column changes, instrument cleaning, or changes of sensitivity of the system over time, a between-batch correction is performed. Whenever more than one batch is present, a batch correction is performed to minimize batch-to-batch variation in the data, by calculating the median ratio of every compound in all the QC samples per batch as described in Equation 2. These batch-to-batch compound-specific correction factors are calculated using Equation 2 and multiplied with the area ratios of each compound, as is shown in Equation 3.

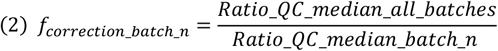

When trends occur within batches, such as a decrease in sensitivity, mzQuality can correct for this using within-batch correction. The within-batch correction is performed using a first-order regression^19^.

#### Compound quality

Once all the batch corrections are performed, various metrics are calculated to check the quality of compounds. The RSDqc is calculated using the QC samples both before and after batch correction, to evaluate the stability of a compound during measurements. Additionally, the percentage of QC samples in which each compound is detected is calculated to improve confidence in the identified compounds. The RSDqc is calculated per compound according to Equation 4 The standard deviation (SD) of the corrected ratios of all QC samples from all batches (SD_corrected_ratio_QC_all_batches) is divided by the mean of all these ratios (Mean_corrected_ratio_QC_all_batches).

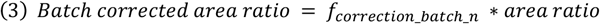

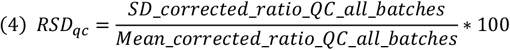

Blank samples are used to assess the background signal. The background signal is calculated according to Equation 5, by dividing the mean of all the areas in all blank samples (Mean_peak area_blanks) by the median of all the areas in study samples (Median_peak area_study samples).

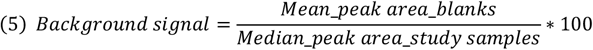

Based on the recommendations of the metabolomics quality assurance and quality control consortium, we set the threshold for the RSDqc at 30%^14^. For the background signal, we recommend that the value should not exceed 40%. However, mzQuality allows users to adjust these settings according to their preferences and analytical methods.

### Data set

The example data set passed the initial integrity and validation tests, and the calculations were performed as described. The following mzQuality output is provided to demonstrate the visualization of data useful in evaluating data quality.

#### Aliquot plot

The initial step of visual assessment of the data is to inspect the aliquot plot per batch. Figure 2 shows the aliquot plot, where the horizontal axis shows the samples in injection order). Panel A shows the median area of all compounds, panel B shows the area_IS and C shows the area ratio, all plotted against the injection order of the samples. This plot portrays all the samples as boxes in a box-whisker plot, color-coded per sample type, and offers insights into the distribution of all measured compounds per sample. In panels A and C, the CAL samples and ACAL samples follow an upward trend, consistent with an expected pattern for a calibration curve with increasing concentrations. The boxplots from QC samples of the same type display similar areas. In panel C, the PROC samples show low values, and the blank samples are high. The ratio of PROC is expected to be low due to the absence of compounds (area) but there are internal standards present. Conversely, the high values in blank samples result from the lack of internal standards. Particularly, the aliquot plot can be used to assess the data before any correction is performed and to check for instrumental drift, missing samples, misinjections, or other mishaps like incomplete batch export from the raw data. Instrumental drift visible in the area (panel A) should not be visible anymore in the ratio plot (panel C). However, extreme instrumental drift may not be corrected away or only affect individual compounds so within-batch correction should then be applied. Figure 2 highlights sample 209 in yellow, showing a notably lower area compared to the other samples in panel A. This could be attributed to either biological or technical issues. Panel C shows the variation of the sample. 209 can be corrected by using IS, making the variation most likely from a technical source and not a biological one.

**Figure 2.**
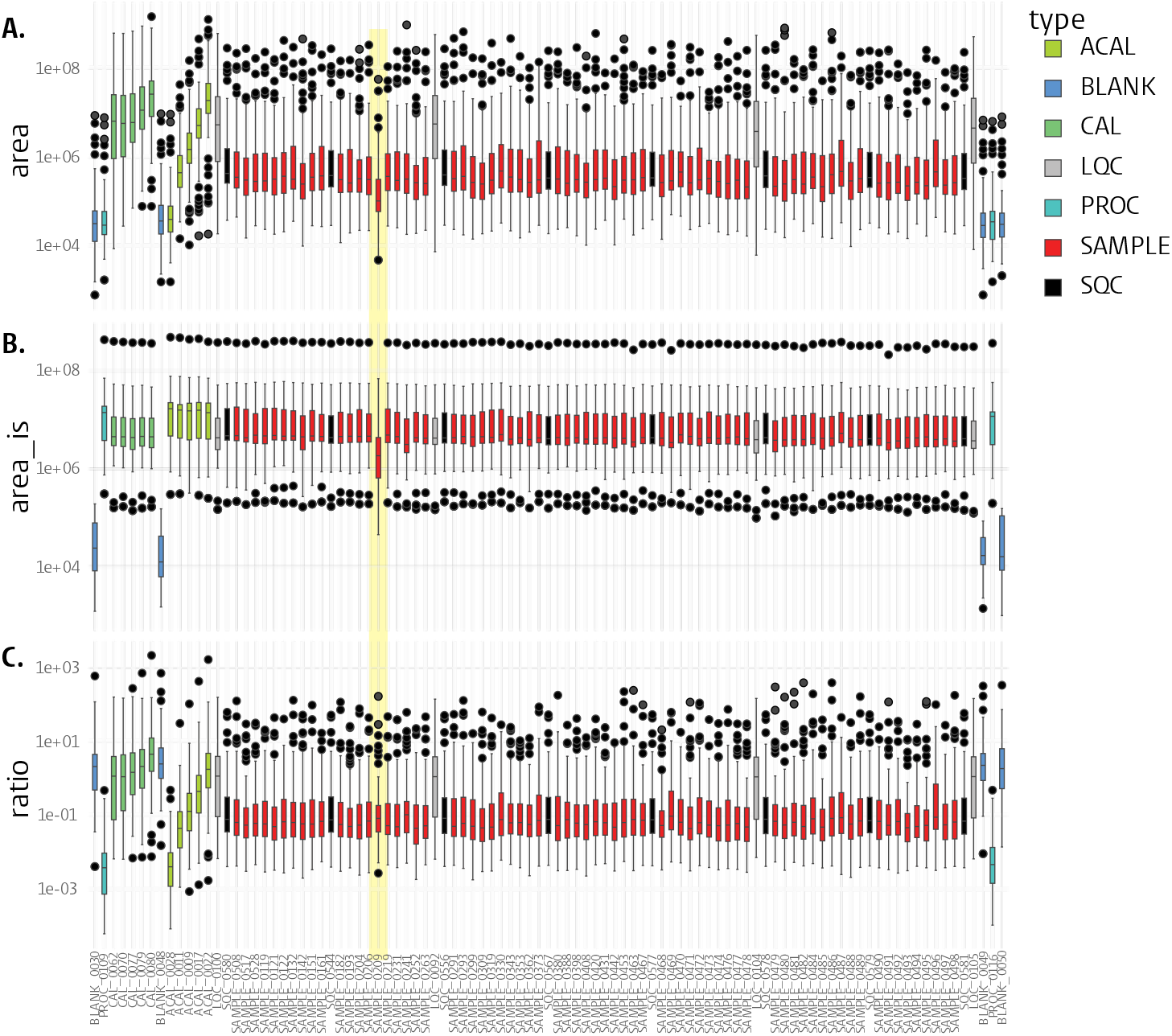
The aliquot plot shows successful internal standard correction of batch one. Every box-whisker plot is color-coded per sample type: calibration mix in water (ACAL; light green), BLANK (dark blue), calibration mix in matrix (CAL; dark green), long-term quality control (LQC; grey), procedure blank (PROC; light blue), SAMPLE (red) and short-term quality control (SQC; black). Black dots represent outlier compounds within the samples. Panel A shows the median of all metabolite areas per sample, panel B shows the median of all the internal standards per sample, and panel C shows the median ratios (area/area_IS) per sample. Sample 0209 is highlighted in yellow due to its relatively low area compared to the other samples.Panel **C**. shows that the variation in sample 0209 is corrected by internal standard use.

#### PCA plot

A PCA plot can be used to 1) check how representative the QC samples are to the study, 2) visualize possible batch effects and 3) identify potential sample outliers. Ideally, SQC samples should be centered as they are aliquots of a pool of all study samples and thus represent their average. Figure 3A shows that before batch correction, the samples and SQC samples from the two batches were not aligned. Still, after batch correction (Figure 3B), the SQC samples are aligned and mostly centered within the study samples, with the two confidence intervals (CI) ellipses overlapping. To reduce the complexity of the PCA plot only two batches are shown while the PCA plots of all batches can be found in Supplementary Figure 1.

**Figure 3:**
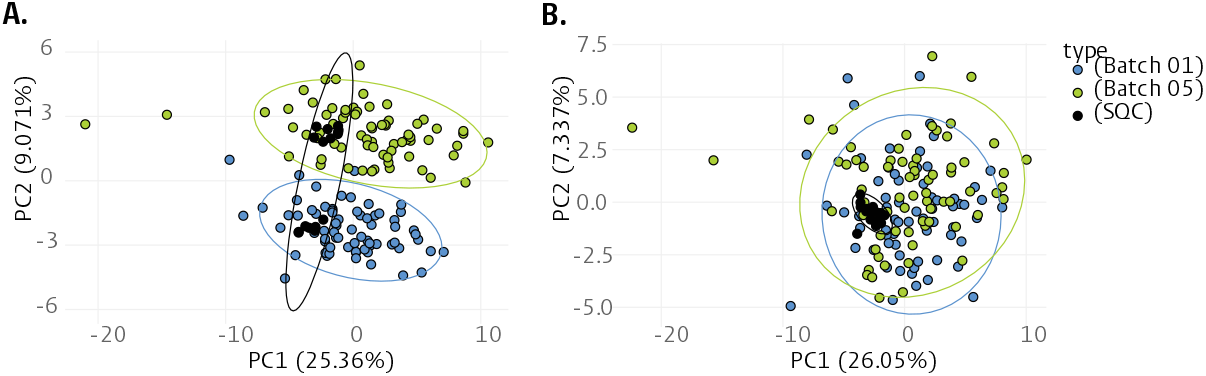
PCA plot before and after between batch correction. **A**. PCA plot of batch 1 and batch 5 before correction. **B**. PCA plot of batch 1 and batch 5 after correction. The samples are grouped by batch and show a 95% confidence interval. After batch correction, the short-term quality control (SQC) samples are aligned in the middle of the two batches.

Outlier samples can be detected by identifying those that fall far outside the 95% CI ellipse. These outliers should be flagged and investigated for deviations in compound peak areas or from standard experimental procedures. If the peaks are normal and no technical explanation is found, the sample should not be removed from the dataset, as the observed variation is most likely due to biological differences.

#### QC violin plot

A violin plot displays QC samples that were not removed in the prior outlined steps, where each dot represents a compound. The plot can be used to investigate QC outliers and deviating batches. Figure 4 illustrates the QC data of three batches before and after between-batch correction. Assessment of the plot involves 1) examining the shape of the violin plots, which should be consistent over the batches, and 2) investigating the median of the QCs which should all align on a horizontal line.

**Figure 4:**
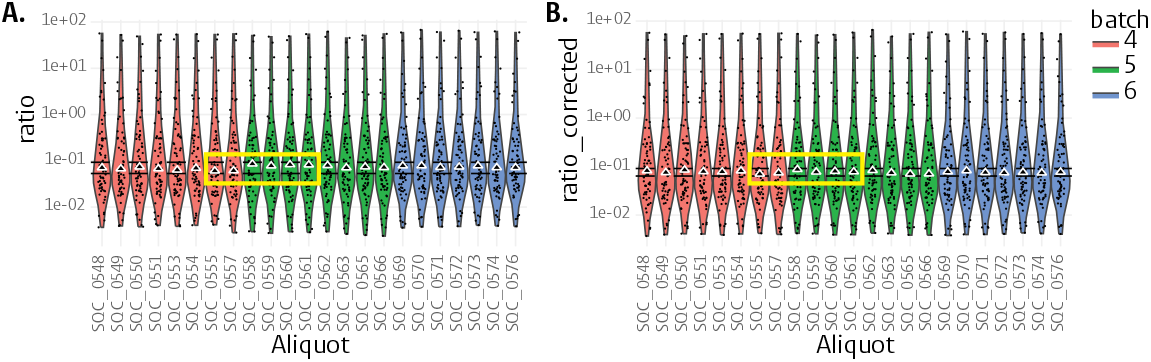
Violin plot of batch 04-06 before and after correction. A. Violin plot of quality control samples from batch 04-06 before batch correction. B. Violin plot of quality control samples from batch 04-06 after batch correction. Short-term quality control (SQC) samples 0555, 0557, 0558, 0559, 0560 and 0561 are highlighted to demonstrate the between-batch correction performed by mzQuality.

Deviations in the median ratio are observed in the last two samples of batch 4 and the first three samples of batch 5 (Figure 4A). Although the sample numbers are not sequential due to the batch design, their measurements are consecutive. The observed deviations are corrected by the between-batch adjustment applied by mzQuality (Figure 4B).

#### Individual compound plots

The individual compound plot can display peak area, area_IS, ratio, corrected ratio, retention time, and other numerical values related to compounds, plotted against the injection number. While compound plots can visualize all sample types, CAL, ACAL, blank, LQC, and PROC are not included in Figure 5 for clarity.

**Figure 5:**
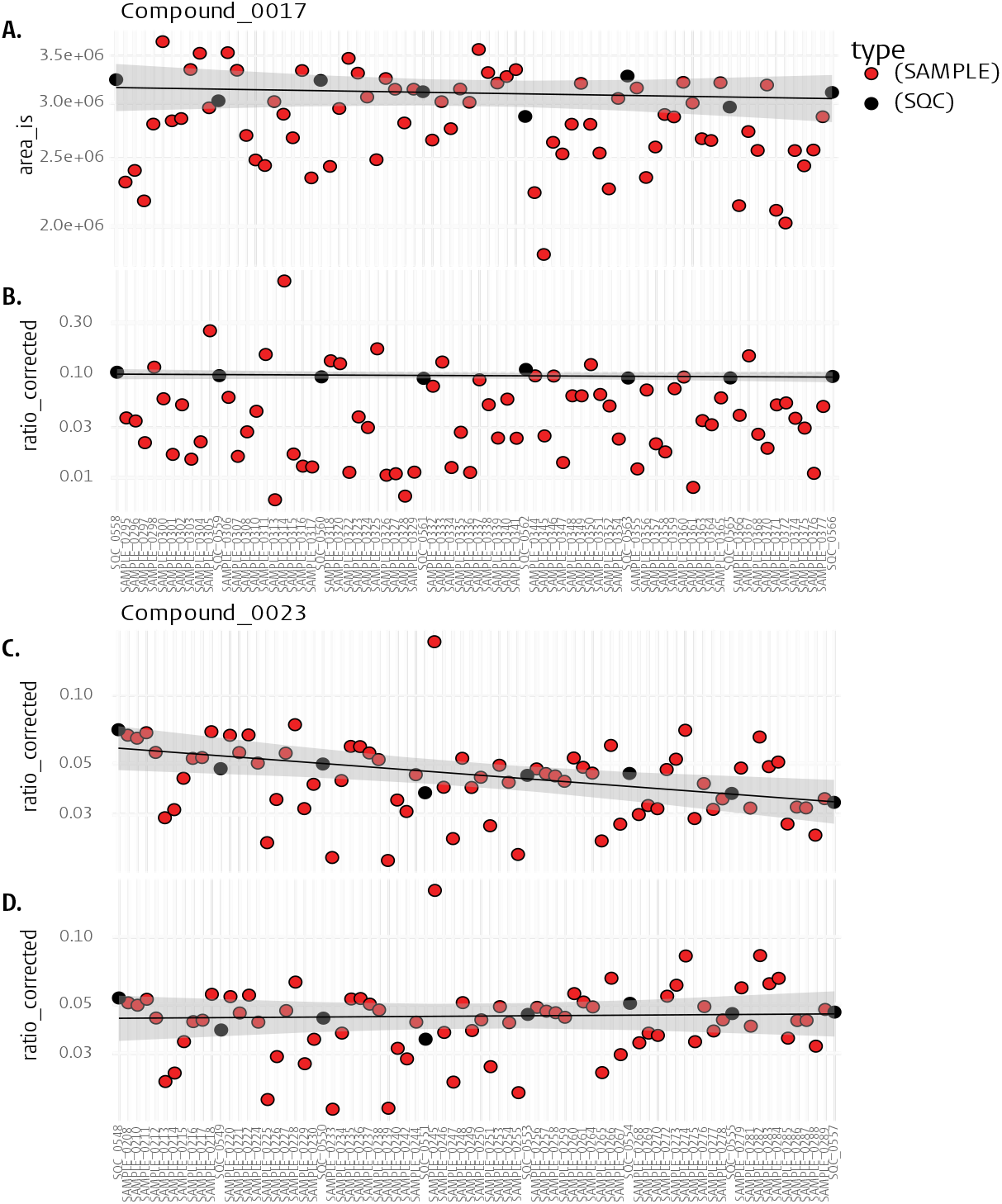
Individual compound plots. the y-axis shows the log_2_ transformed corrected ratios, while the x-axis shows the injection number. Red dots portray the samples, while black dots portray the short-term quality control (SQC)**A**. shows the area_IS of the internal standard related to compound 17 in batch 5. **B**. shows an example of a compound without any trends in batch 5. **C**. illustrates an example of a downward trend within one batch, and **D**. shows how this trend is corrected with the within-batch correction functionality of mzQuality both in batch 4.

Figure 5A displays an individual compound plot for an IS. These plots provide a unique ability to visually inspect the IS quality of individual samples within mzQuality.

Figure 5B provides an example of a compound plot without trends over time or other irregularities. Here, the between-batch corrected area ratio is plotted against the injection number of samples which can reveal trends within and across batches.

Figure 5B and Figure 5C demonstrate the effect of within-batch correction where before (Figure 5B) a noticeable downward trend over time is observed which is no longer present in Figure 5C. Within-batch correction is applied to all compounds, so its effects should be checked for each compound. The additional plots including all sample types can be found in Supplementary Figure 2. These additional plots can be used to monitor trends in retention time. This is particularly valuable for identifying potential data integration errors. Samples with missing values, as indicated by NA, will not have a data point shown.

After visually inspecting all individual compound plots, it may be necessary to revisit peak integration to address any observed irregularities. If these irregularities can be corrected, new export files should be generated, and mzQuality must be run from the beginning. This output folder includes all the previously mentioned plots and Excel files that can be used for statistical analysis in tools such as R.

The Excel file lists all measured compounds, categorized by confidence level. Compounds are organized into three tabs based on quality and user-defined or default settings the ‘High confidence’ tab includes compounds with RSDqc values below 15%, the ‘With Caution’ tab includes compounds with RSDqc values ranging between 15% and 30%, and the ‘Low Signal to Noise’ tab including compounds with either background signal >40% or with RSDqc values > 30%. An example and further details on the content in the output folder are provided in GitHub: https://github.com/hankemeierlab/mzQuality

## Conclusions

Here, we outlined practices to minimize experimental variation and to ensure optimal performance of mzQuality. Additionally, we showcased a typical application of mzQuality with a supplied dataset, collected across multiple batches. The software can manage any type of raw data, independent of vendor software, and does not require the user to have any programming skills. Its intuitive visualization of data enables users to assess their data easily. Moreover, mzQuality assures the user that the data is suitable for statistical and biological interpretation in line with the requirements set by regulatory bodies like the FDA and EMA. By adhering to these recommended practices and by utilizing mzQuality, we anticipate that the metabolomics community will generate higher-quality data, thereby improving the quality of data reported in the literature.

## Funding

Co-funded by the European Union’s Horizon Europe Framework Programme (101080997, Recon4IMD)

Co-funded by the CHDR as part of the SKINERGY trial under NWA-ORC project NWA.1389.20.182 entitled Next Generation ImmunoDermatology (NGID).

Co-funded by X-Omics (NWO, project 184.034.019) The research is supported by the NWO Gravitation programme EXPOSOME-NL (grant number 024.004.017).

## Notes

https://github.com/hankemeierlab/mzQuality

### Competing Interest Statement

The authors have declared no competing interest.

## Reference

1. Chu, X. et al. Integration of metabolomics, genomics, and immune phenotypes reveals the causal roles of metabolites in disease. Genome Biol. 22, 198 (2021).

2. Kohler, I., Hankemeier, T., van der Graaf, P. H., Knibbe, C. A. J. & van Hasselt, J. G. C. Integrating clinical metabolomics-based biomarker discovery and clinical pharmacology to enable precision medicine. Eur. J. Pharm. Sci. 109, S15–S21 (2017).

3. Allard, P.-M. et al. Integration of Molecular Networking and In-Silico MS/MS Fragmentation for Natural Products Dereplication. Anal. Chem. 88, 3317–3323 (2016).

4. Han, W. & Li, L. Evaluating and minimizing batch effects in metabolomics. Mass Spectrom. Rev. 41, 421–442 (2022).

5. Livera, A. M. D. et al. Statistical Methods for Handling Unwanted Variation in Metabolomics Data. Anal. Chem. 87, 3606–3615 (2015).

6. Liu, Q. et al. Addressing the batch effect issue for LC/MS metabolomics data in data preprocessing. Sci. Rep. 10, 13856 (2020).

7. Wehrens, R. et al. Improved batch correction in untargeted MS-based metabolomics. Metabolomics 12, 88 (2016).

8. Broadhurst, D. et al. Guidelines and considerations for the use of system suitability and quality control samples in mass spectrometry assays applied in untargeted clinical metabolomic studies. Metabolomics 14, 72 (2018).

9. Brown, M. et al. A metabolome pipeline: from concept to data to knowledge. Metabolomics 1, 39– 51 (2005).

10. Drotleff, B. & Lämmerhofer, M. Guidelines for Selection of Internal Standard-Based Normalization Strategies in Untargeted Lipidomic Profiling by LC-HR-MS/MS. Anal. Chem. 91, 9836–9843 (2019).

11. Lange, E., Gröpl, C., Reinert, K., Kohlbacher, O. & Hildebrandt, A. High-accuracy peak picking of proteomics data using wavelet techniques. Pac. Symp. Biocomput. Pac. Symp. Biocomput. 243–254 (2006).

12. MZmine 2 - Features. https://mzmine.github.io/features.html#Batch.

13. Naake, T., Rainer, J. & Huber, W. MsQuality: an interoperable open-source package for the calculation of standardized quality metrics of mass spectrometry data. Bioinformatics 39, btad618 (2023).

14. Kirwan, J. A. et al. Quality assurance and quality control reporting in untargeted metabolic phenotyping: mQACC recommendations for analytical quality management. Metabolomics 18, 70 (2022).

15. Dunn, W. B. et al. Quality assurance and quality control processes: summary of a metabolomics community questionnaire. Metabolomics 13, 50 (2017).

16. Yang, W. et al. A comprehensive UHPLC-MS/MS method for metabolomics profiling of signaling lipids: Markers of oxidative stress, immunity and inflammation. Anal. Chim. Acta 1297, 342348 (2024).

17. Baggerly, K. A., Edmonson, S. R., Morris, J. S. & Coombes, K. R. High-resolution serum proteomic patterns for ovarian cancer detection. Endocr. Relat. Cancer 11, 583–584 (2004).

18. R: Rosner’s Test for Outliers. https://search.r-project.org/CRAN/refmans/EnvStats/html/rosnerTest.html.

19. van der Kloet, F. M., Bobeldijk, I., Verheij, E. R. & Jellema, R. H. Analytical error reduction using single point calibration for accurate and precise metabolomic phenotyping. J. Proteome Res. 8, 5132–5141 (2009).

