## Supplementary Figures for "mzQuality: A tool for quality monitoring and reporting of targeted mass spectrometry measurements"

**A.**

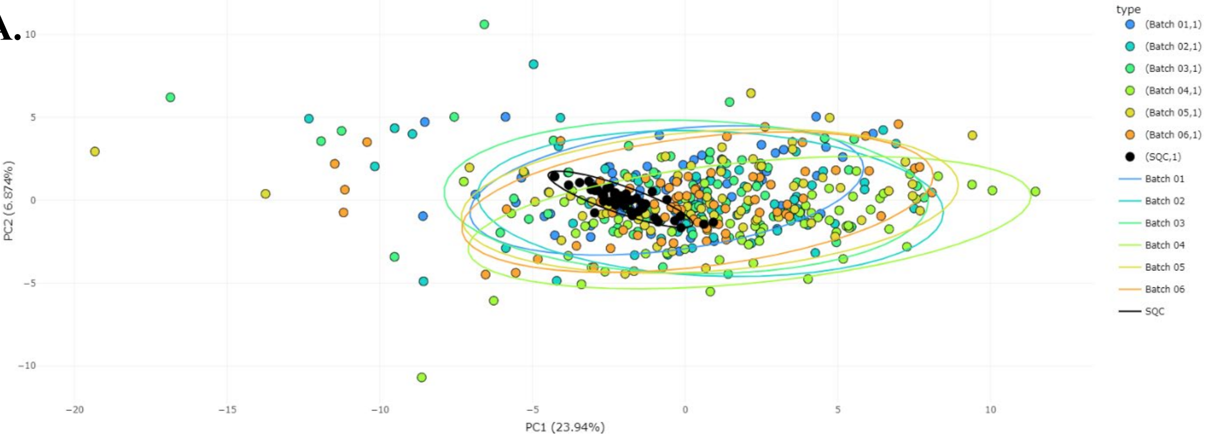


**B.**

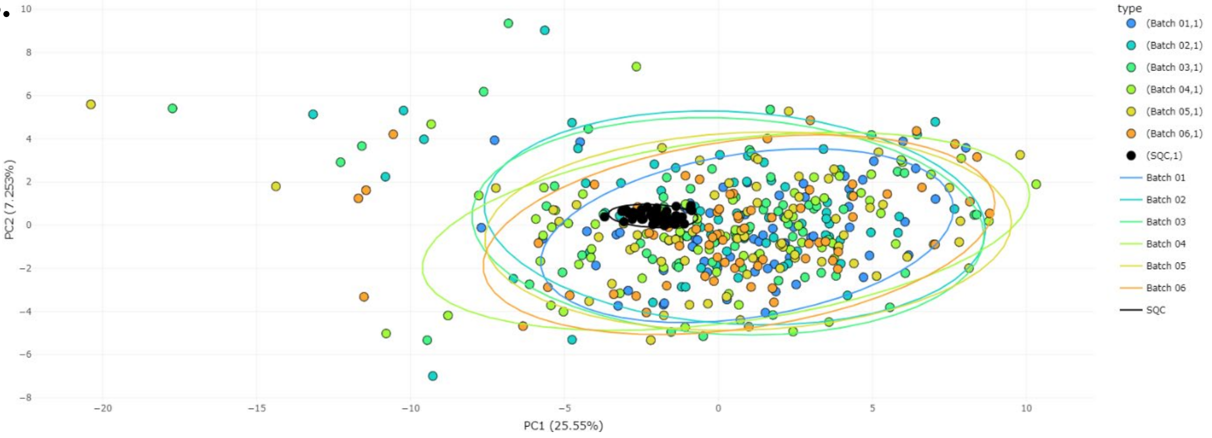


**Supplementary Figure 1: PCA plot before and after between-batch correction**. **A**. PCA plot of batch 1-6 before correction. **B**. PCA plot of batch 1-6 after correction. The samples are grouped by batch and show a 95% confidence interval. After batch correction, the SQC samples are more aligned in the middle of the two batches.

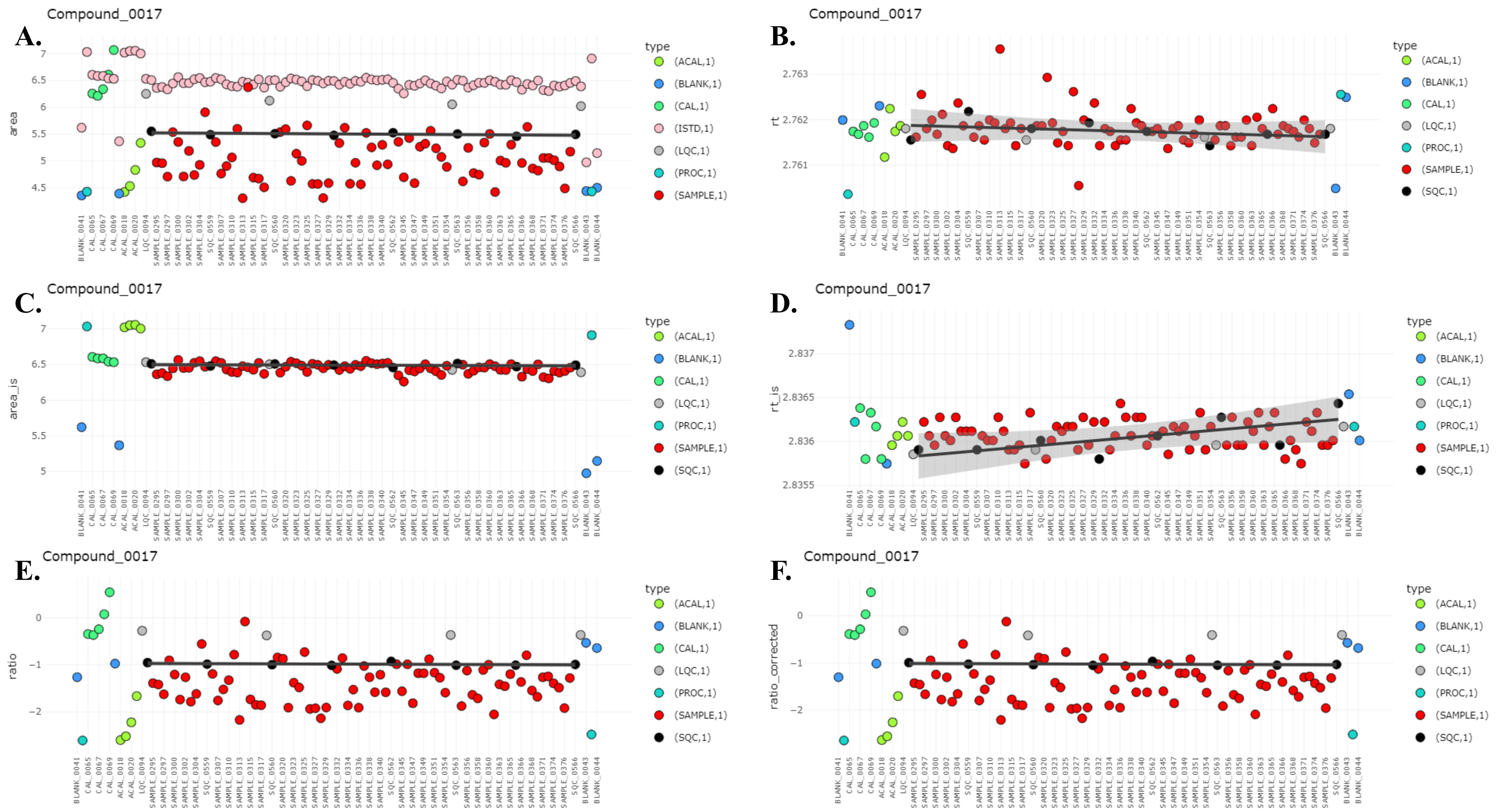


**Supplementary Figure 2: Individual plots of compound 17 in batch 5 including all sample types.** The y-axis displays the log2 transformed corrected ratios, and the x-axis displays the injection number. **A**. Area including the internal standard area. **B.** Retention time **C**. Area of the corresponding internal standard **D**. Retention time of the internal standard (in minutes). **E**. Ratio of the compound area to the internal standard area (area/area_is). **F.** Corrected ratio after between-batch correction.
